# A genetic toolkit for investigating *Clavibacter*: markerless deletion, permissive site identification and an integrative plasmid

**DOI:** 10.1101/2021.07.13.452269

**Authors:** Danielle M. Stevens, Andrea Tang, Gitta Coaker

## Abstract

The development of knockout mutants and expression variants are critical for understanding genotype-phenotype relationships. However, advancements of these techniques in Gram-positive actinobacteria have stagnated over the last decade. Actinobacteria in the *Clavibacter* genus are composed of diverse crop pathogens which cause a variety of wilt and cankering diseases. Here, we present a suite of tools for genetic manipulation in the tomato pathogen *C. michiganensis* including a markerless deletion system, an integrative plasmid, and an R package for identification of permissive sites for plasmid integration. The vector pSelAct-KO is a recombination based, markerless knockout system that uses dual selection to engineer seamless deletions of a region of interest, providing opportunities for repeated higher-order genetic knockouts. The efficacy of pSelAct-KO was demonstrated in *C. michiganensis* and confirmed using whole genome sequencing. We developed permissR, an R package to identify permissive sites for chromosomal integration, which can be used in conjunction with pSelAct-Express, a non-replicating integrative plasmid that enables recombination into a permissive genomic location. Expression of eGFP by pSelAct-Express was verified in two candidate permissive regions predicted by permissR in *C. michiganensis*. These molecular tools are essential advancements for investigating Gram-positive actinobacteria, particularly for important pathogens in the *Clavibacter* genus.

## Introduction

Gram-positive actinobacteria in the *Clavibacter* genus are economically important xylem-colonizing bacterial pathogens that can infect both monocots and dicots, resulting in canker and wilting diseases (Eichenlaub and Gartemann 2011; Thapa et al. 2019). Despite current disease control measures including good horticultural practices and monitoring seed stock, outbreaks of *Clavibacter* pathogens have occurred in recent years (Peritore-Galve et al. 2021). The corn pathogen *C. nebraskensis* has been problematic for many corn-producing Midwestern states and the tomato pathogen *C. michiganensis* has caused notable losses in years past (Ahmad et al. 2018; Nandi et al. 2018; Peritore-Galve et al. 2021). While outbreaks are sporadic, four of the six species are considered quarantine organisms by the European and Mediterranean Plant Protection Organization with *C. michiganensis* acting as a potential threat to both greenhouse and field production (Nandi et al. 2018; Peritore-Galve et al. 2021). Despite their agricultural impact, little is known about how these pathogens cause disease (Nandi et al. 2018; Thapa et al. 2019).

In other bacterial pathogens, genes important for pathogenicity and host range include secreted protein effectors that suppress host immunity, alter host metabolism, and enable colonization, providing a fitness advantage. The most well studied *Clavibacter* species is the tomato pathogen *C. michiganensis. C. michiganensis* carries effectors on two plasmids, pCM1 and pCM2, and within the ~129 Kb *chp/tomA* pathogenicity island (PAI) (Meletzus et al. 1993). While the entire *chp/tomA* PAI is only found in *C. michiganensis*, fragments and/or homologs of PAI members can be found in the genomes of other pathogenic *Clavibacter* species (Bentley et al. 2008; Tambong 2017; Hwang et al. 2018; Lu et al. 2018). The chp/tomA PAI encodes suites of putative serine proteases belonging to the Sbt, Chp, and Ppa families and carbohydrate activating enzymes (CAZymes) (Thapa et al. 2017). A few *C. michiganensis* effectors have been characterized, including the tomatinase, *tomA* and the cellulase, *celA* (Meletzus et al. 1993; Jahr et al. 2000). However, the plant targets of most effectors are unknown (Nandi et al. 2018).

Pioneering work in the late 1990s through early 2010s resulted in vectors for gene deletion and expression in *Clavibacter*, but there is still reliance on these tools despite known limitations. Mutants depend on gene replacement with an antibiotic cassette or by transposon mutagenesis (Kirchner et al 2001; Peritore-Galve et al. 2021). Engineering gene expression relies on using a plasmid modified from the backbone of native pCM1/pCM2 plasmids from *C. michiganensis*. This requires recipient strains to lack the native plasmid, a confounding factor because known virulence genes are carried on pCM1/pCM2. Gene expression can also be achieved by transposition via a transposase-based vector, which can have unexpected effects depending on the genetic location inserted (Laine et al. 1996; Chalupowicz et al. 2012; Tancos et al. 2013).

Genetic tool kits in other bacterial systems enabled functional characterization of genetic drivers in disease development. The original pSelAct vector was designed to make gene deletions in the foal actinobacterial pathogen, *Rhodococcus equi* (Van der Geize et al. 2008). pSelAct was modified through the addition of a Gateway cassette into the multiple cloning site and used to make single gene deletions in the distantly related actinobacterial plant pathogen, *Rhodococcus fascians* (later referred to as pSelAct-KO; Savory et al. 2020). A second group independently developed pMP201 for deletions in *Streptomyces*, which functions similarly to the pSelAct vector using a positive selection marker and counter selection (Dubeau et al. 2008). In *Ralstonia solanacearum* GMI1000 and related strains, the integrative plasmid series pRC enables insertion and expression of genetic material in a defined location (Monteiro et al. 2012). Finally, a knock-out vector using the sucrose counter selection marker, *sacB*, was modified for integrative expression in *Pseudomonas syringae* pv. *tomato* DC3000 (Lee et al. 2018). Flexible genetic tools facilitate rapid advances in studying pathogens regardless of the system.

Here, we present a suite of genetic tools to facilitate investigation of *Clavibacter*, including a markerless deletion system, an integrative plasmid, and an R package for identification of permissive sites for plasmid integration. We optimized pSelAct-KO, a recombination-based system that allows for development of higher order markerless deletions. pSelAct-KO was used to make a 5.6 kB deletion in the *C. michiganensis chp/tomA* PAI comprising five effectors (*chpE* through *ppaC*) and the deletion was confirmed using whole genome sequencing. We developed the R package, permissR, and pSelAct-Express to work in conjunction to allow for identification of permissive genomic locations and subsequent targeted plasmid integration. In total, these tools should accelerate reverse genetics and subsequent genotype-phenotype studies in *Clavibacter*.

## Results and Discussion

### pSelAct-KO system for markerless deletions in *Clavibacter*

The utility of two-step systems has been demonstrated in other actinobacteria including *Rhodococcus* and *Streptomyces* bacteria as well as other Gram-positive bacteria (Dubeau et al. 2008; Kostner et al. 2017; Chen et al. 2020; Savory et al. 2020). Thus, generating markerless deletions using selection and counterselective methods could likely be applied *Clavibacter* and other related taxa. We employed the pSelAct-KO vector to *Clavibacter* with the strategy for construct design, selection, and counterselection in Figure 1. In order to develop the knockout construct, approximately 1.5 kilobase (kb) regions flanking the gene/gene locus of interest are amplified (Fig. 1A). Fragments from the 5’ and 3’ flanking regions are cloned into the pSelAct-KO Gateway C Cassette, through either Gateway cloning or a homology-based cloning approach (In-Fusion or Gibson) to generate pSelActΔ*goi* (Fig. 1B) (Savory et al. 2020). pSelActΔ*goi* is transformed into *E. coli*, screened via PCR, and verified via Sanger sequencing, then subsequently transformed into the wildtype *Clavibacter* strain (Fig. 1C). Positive recombinants are resistant to the antibiotic apramycin and are screened for their merodiploid state (haploid cells which have duplicated genetic material) using PCR (Fig. 1D). Growing the merodiploid on minimal medium supplemented with 5-fluorocytosine (5-FC) will promote a recombination event to generate a clean (scarless) deletion (Fig. 1E).

**Figure 1:**
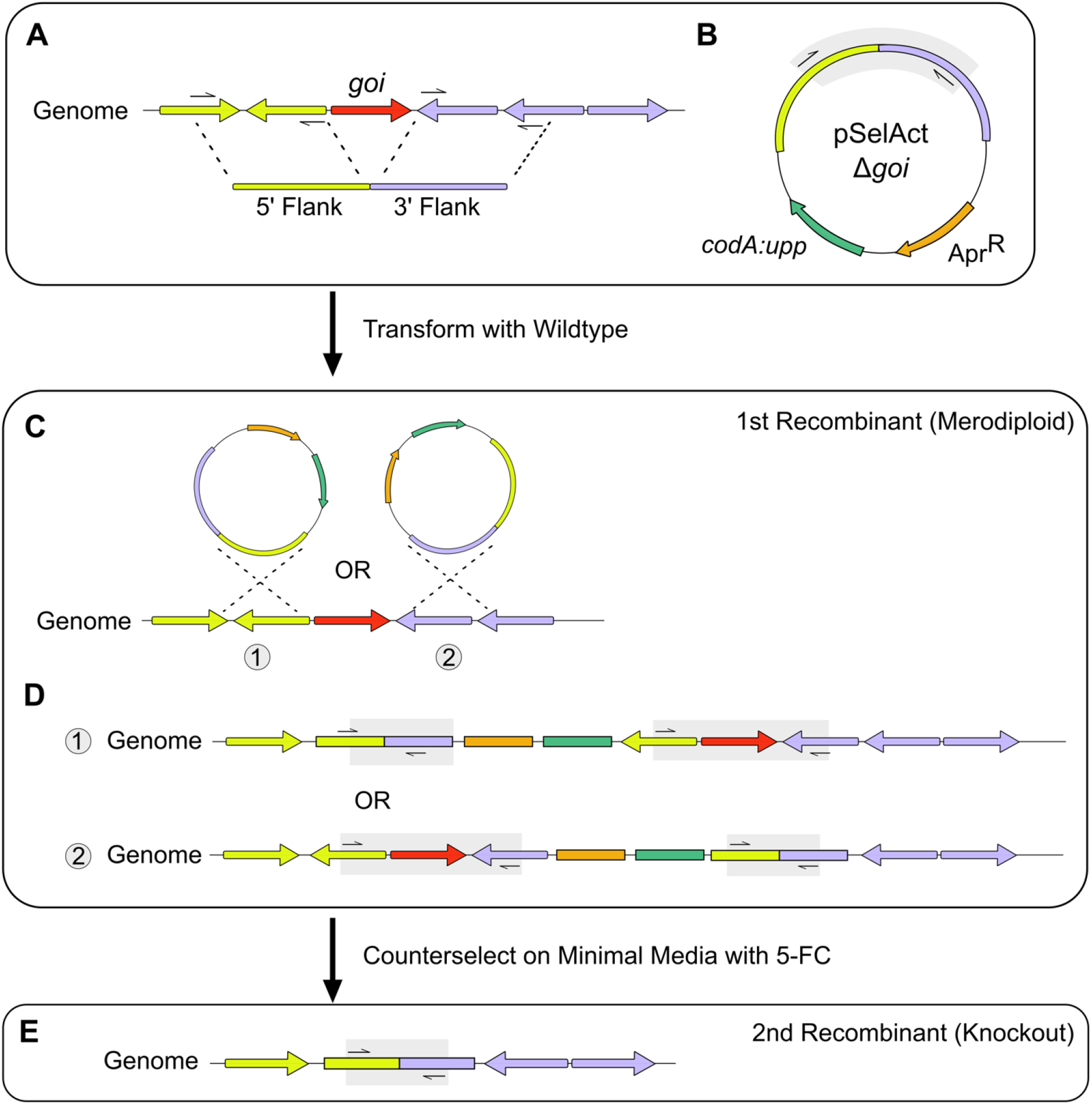
Pipeline for markerless knockouts using pSelAct-KO. Grey shaded area signifies the amplified region using primers spanning the region of interest. **A.** Amplification of flanking regions before and after gene(s) of interest. **B.** Cloning of 5’ and 3’ flanking regions into pSelAct-KO vector through either Gateway or homology-based cloning. **C.** Possible sites of recombination during transformation and selection of non-replicating vector into the genome. **D.** Possible merodiploid gene structure. **E.** Gene structure after recombination where the region of interest is deleted.

#### Determining apramycin resistance in C. michiganensis

The pSelAct-KO system employs selection using the aminoglycoside antibiotic apramycin to select for chromosomal integration followed by counterselection with 5-FC to recombine out of the genome. To determine the minimum inhibitory concentration (MIC) of apramycin, MIC growth curves were performed in *C. michiganensis*. In liquid rich tryptone broth with yeast (TBY) media, the MIC necessary to limit growth was 50 μg/mL (Fig. 2B). On solid agar TBY plates, the MIC required was higher at 100 μg/mL (Supplemental Fig. S1). Notably, if plates were incubated beyond five days post *C. michiganensis* transformation, smaller colonies appeared to form (data not shown). This appeared to dissipate if *Clavibacter* transformants were plated on TBY that had double the MIC at 200 μg/mL. However, TBY plates with 100 μg/mL apramycin were sufficient for downstream screening (Supplemental Table S1). Susceptibility to the antibiotic apramycin in *C. michiganensis* differed on plates compared to other actinobacterial pathogens, which require lower concentrations (Savory et al. 2020). A TblastN search of the apramycin resistance gene in *Clavibacter* (taxid: 1573) yielded no significant results and antibiotics within the same drug class, neomycin and gentamycin, are known to be effective (Supplemental Table S2) (Meletzus and Eichenlaub 1991). While it is unclear why a higher concentration is required, apramycin is affective against *C. michiganensis*.

**Figure 2:**
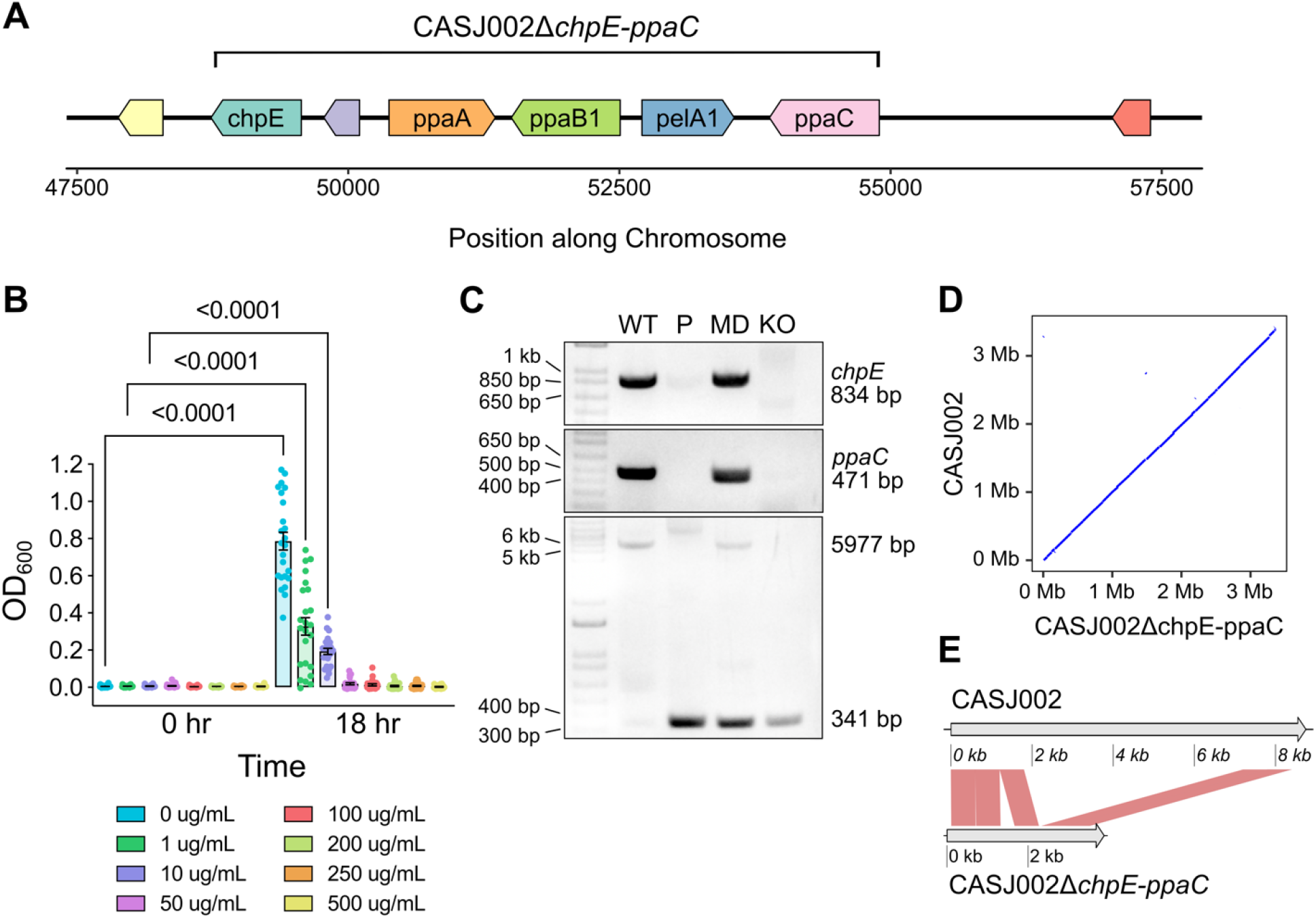
Markerless deletion of *chpE* through *ppaC* in the tomato pathogen *C. michiganensis* CASJ002. **A.** Map of genes of interest for deletion and their flanking regions. **B.** Minimum inhibitory concentration of the antibiotic apramycin in TBY liquid culture for *C. michiganensis* isolate CASJ002. Eight replicates per treatment were performed with media only as a negative control. The experiment was repeated independently three times. A one-way ANOVA and a post-means Tukey’s multiple comparisons test was computed. Comparisons of 0 hr and 18hr values from 50 ug/mL upward were not significantly different. **C.** PCR-based verification of deletion on 1% TAE agarose gel. Top: Gene specific primers for *chpE*. Middle: Gene specific primers for *ppaC*. Bottom: Primers which bind to the flanking regions and span the region deleted. Lanes are products from the reactions including: WT; Genomic DNA from CASJ002, P: Plasmid DNA from *E. coli* carrying the pSelAct_CASJ002_chpE-ppaC_KO; MD: Genomic DNA from the merodiploid; KO: Genomic DNA from CASJ002Δ*chpE-ppaC*. The estimated sizes of the PCR products are depicted (bp = base pairs). **D.** Dot plot of CASJ002Δ*chpE-ppaC* against reference CASJ002 genome sequence. **E.** Syntenic comparison of the region of interest from CASJ002 against the engineered deletion in CASJ002Δ*chpE-ppaC*. Comparison fragment size is 600 bp. The gap in the 5’Flanking region of CASJ002Δ*chpE-ppaC* represents a gap between the scaffolded contigs.

#### Deletion of 5.6kB gene cluster in the tomato pathogen C. michiganensis

To demonstrate the utility of the pSelAct-KO for gene deletion, we focused on the *chp* region in the PAI of *C. michiganensis* strain CASJ002. PAI-localized effectors have been of interest since their discovery and are hypothesized to drive disease development in the *C. michiganensis*-tomato pathosystem (Eichenlaub and Gartemann 2011; Nandi et al. 2018; Thapa et al. 2019). Within the *chpE-ppaC* region, there are several known virulence genes, including four serine proteases and the CAZyme *pelA1* (Fig. 2A). Three proteases are from the Ppa family and one from the Chp family. Four of the five genes of interest have been individually deleted (Chalupoqicz et al. 2017; Thapa et al. 2017). Mutants of the proteases *chpE, ppaA*, and *ppaC* had no reported effect on disease development when tested on tomato (Chalupoqicz et al. 2017). The CAZyme *pelA1* mutant had reduced wilting symptoms on tomato (Thapa et al. 2017). The protease *ppaB1* has yet been tested for its contribution in disease development.

To test pSelAct-KO in *C. michiganensis*, we developed primers to amplify flanking regions before *chpE* and after *ppaC* (Fig. 2A). Using homology-based In-Fusion cloning, the fragments replaced the Gateway cassette of pSelAct-KO and a positive clone was selected and sequence verified (Fig. 1A and B). Upon transformation of *C. michiganensis* wildtype strain CASJ002, the vector will recombine with one of the two homology arms (Fig. 1C). PCR was used to screen for the merodiploid state; first recombinants will provide two bands: one that is the same as wildtype and one that is smaller, representative of the recombination event (Fig. 1D: grey box highlights two amplicons; Fig. 2C). It is important to note that as the deletion size increases, it may be more difficult to amplify the upper band representing the wildtype state.

To develop the knockout through secondary recombination, the merodiploid was plated in dilutions from 10^-6^ to 10^-7^ on minimal M9 medium supplemented with the counterselectant 5-FC. Strains are required to be insensitive to the cysteine analog 5-FC but sensitive to its more toxic counterpart 5-florouracil (5-FU) and 5-fUMP (Debeau et al. 2008). For counterselection, the gene cassette *codA::upp* converts 5-FC into the toxic compound 5-FU and then subsequently 5-fUMP (van der Geize et al. 2008). After 7-14 days of growth, single colonies are visible. As a quick and easy screen, colonies from M9 5-FC plates were re-streaked on both TBY and apramycin-containing TBY plates. Recombination of the vector will inhibit growth on apramycin. Therefore, colonies growing on TBY but not TBY with apramycin either reverted back to wildtype or are a true knockout (Fig. 1E). After incubation at 28°C on M9 5-FC plates, several hundred colonies from the *C. michiganensis* CASJ002 *chpE-ppaC* merodiploid were screened for second recombinants after transferring to apramycin-containing TBY plates. All colonies remained merodiploid, likely showing ineffective compound uptake.

We hypothesize that rapid growth could signal inefficient uptake of 5-FC leading to a non-selective environment. Therefore, adjustments to the incubation temperature were made. We grew the *C. michiganensis* CASJ002 *chpE-ppaC* merodiploid at 24°C on M9 5-FC plates, which results in slower colony growth. Of the 63 colonies subsequently screened on both TBY and apramycin-containing TBY plates at 28°C, three were confirmed to be true knockouts (Supplemental Fig. S2). Previous deletions using pSelAct or other similar systems tend to delete one to two genes at a time, so while we do not know the upper limits of the system, we were able to make a deletion as large as 5,636 bp in *C. michiganensis* CASJ002 (Fig. 2) (Savory et al. 2020). These results highlight the importance of environment conditions impacting 5-FC uptake and effectiveness as means for counterselection.

Admittedly, converting the merodiploid to a knockout mutant can be the most laborious and unpredictable step of the system. However, there are known approaches that can enhance recombination efficiency. One, the concentration of 5-FC could be optimized. We did not choose to alter this concentration based on its success in distantly related actinobacteria and early reports of 5-FC resistance, 5-FU susceptibility in *C. sepedonicus* (Savory et al. 2020; Syverson 2011). Two, a similar system in *Bacillus llicheniformis*, pKVM4, noted that the addition of *codB*, an encoded cytosine permease, notably increased efficiency of selection (Kostner et al. 2017). Koster et al (2017) hypothesized that the addition of *codB* increased 5-FC uptake leading to an increase in the concentration of 5-FU and 5-fUMP present and thus, greater selection. Re-engineering the pSelAct-KO to include a cytosine permease homolog may improve recombination efficiency. Finally, modify environmental conditions can have dramatic effects. In the original pSelAct vector developed for *R. equi*, growing the merodiploid on rich medium over minimal medium demolished the counterselective effects from 5-FC, likely due to a lack of 5-FC uptake related to the pyrimidine salvage pathway. While it is unclear whether decreasing the temperature had the same effect, it is clear environmental conditions can impact recombination frequency.

Collectively, these results demonstrate that pSelAct-KO can be used in *C. michiganensis* to generate markerless deletions. Furthermore, optimizing the growth temperature when the bacteria are grown on M9 with 5-FC resulted in relatively high counterselection. It is important to note that pSelAct-KO has been used in other Gram-positive actinomycetes. Therefore, it is likely that this system will be functional in other *Clavibacter* species and related genera.

#### Sequence confirmation of C. michiganensis CASJ002ΔchpE-ppaC reveal no major structural changes

The strong selection pressure from 5-FC compound conversion to 5-FU and 5-fUMP may result in plasmid loss or other structural changes. Plasmid loss, particularly pCM2, has been noted in past studies that genetically altered *C. michiganensis* (Stork et al. 2008). Of the three confirmed *C. michiganensis* CASJ002Δ*chpE-ppaC*, only one lost the native plasmid, pCM2 (Supplemental Fig. S2). In order to investigate for any potential large structural rearrangements, Illumina paired end reads of *C. michiganensis* CASJ002Δ*chpE-ppaC* colony 9 were assembled and compared to the wildtype genome (detailed methods can be found in Material and Methods as well as in the GitHub repository – pSelAct_KO_Clavibacter). Even with low 13x Illumina sequencing coverage, most contigs were syntenic (Fig. 2D). A clean deletion in the region of interest was further confirmed between the wildtype and deletion (Fig. 2E). Therefore, we recommend screening via PCR for native plasmid(s) as a background assessment and if possible, low coverage whole genome Illumina sequencing to compare against the wildtype genome as a precautionary measure. Since this system is markerless, higher order deletions can be made, showcasing the potential strength of the system especially for characterizing redundant virulence genes or effectors.

### Prediction of permissive sites via permissR

In addition to establishing a system for markerless deletions in *Clavibacter*, we sought to establish an integrative plasmid for gene expression. Unlike previous integrative plasmids which were designed with a specific genome in mind, we wanted to design a system that was flexible for future genomic-driven functional studies. This required developing the necessary tools to predict potential permissive, non-coding target regions for plasmid integration.

We developed an R package, permissR, to predict permissive sites for potential plasmid integration using draft genomes. permissR minimally requires two input files, a GenBank file (.gbff or .gbk format) and a whole genome Fasta file (Supplemental Fig. S3). We also recommend running ISEScan to avoid cloning and recombining into an IS element that could cause unwanted structural changes. IS element prediction can be ran within the R console or separately on the terminal and imported into permissR. Upon running permissR, either by providing paths to the required files or using the pop-up GUI to select the files, the pipeline will provide several outputs. Outputs include a whole genome plot with information on the annotation, IS elements (if present), a scanning 1000-bp window of Shannon entropy and GC-content, two different measurements of complexity (Fig. 3A) (Akhter et al. 2013). Predicted candidate permissive sites are at least 1.5 kb in size, not within IS elements, and located on contigs a minimum 15 kB in size. Plots of candidate sites are outputted such that the user can refer to the neighboring genes (Fig. 4B). Predicted sites are ranked and suppled to the user as a tab-delimited text file. Additionally, candidate sites are also compared against the whole genome and if close hits are present (i.e. the maximum number of mismatches equal 15% of less difference based on permissive hit length), they are exported in a separate tab-delimited text file.

**Figure 3:**
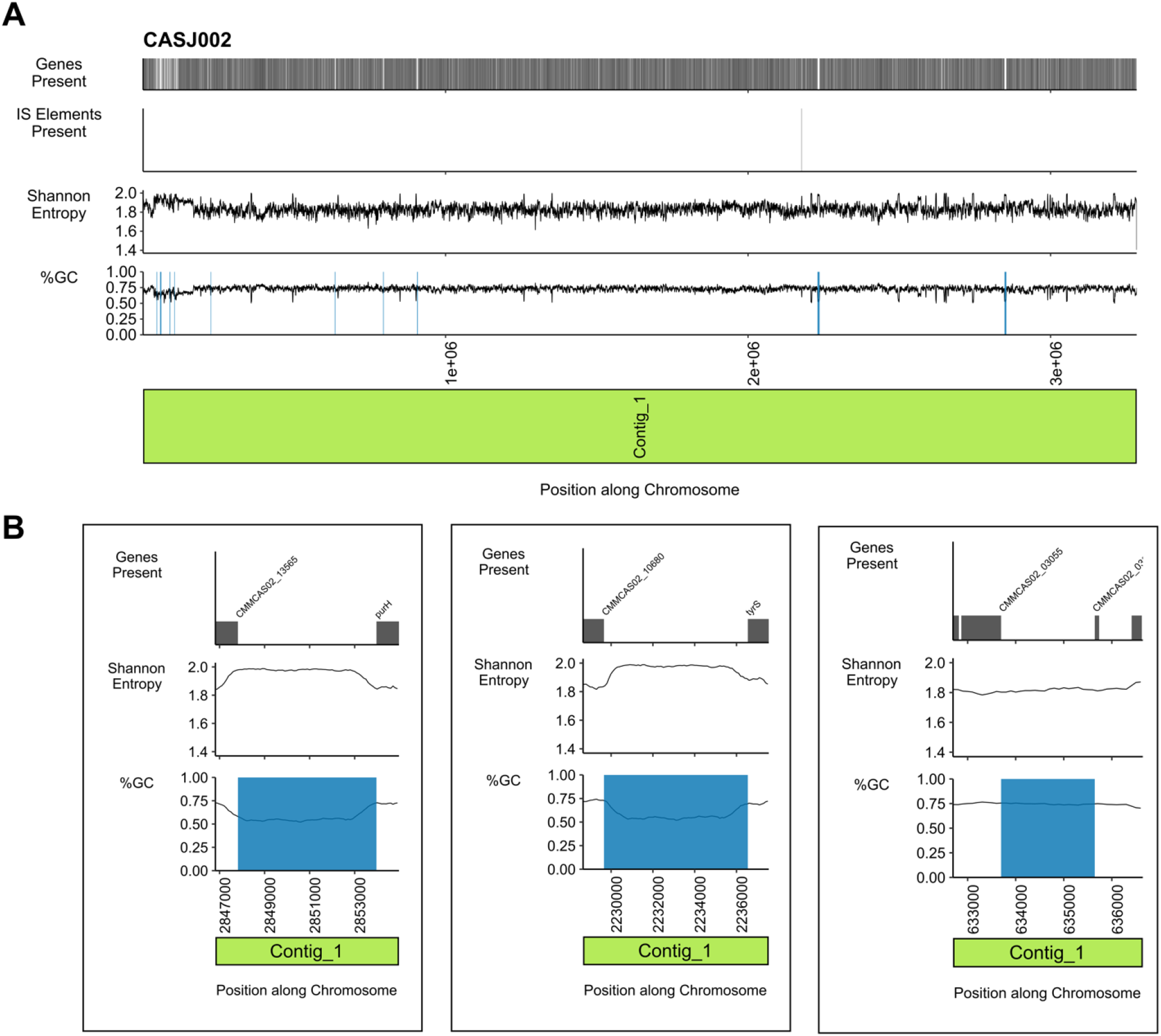
The R package permissR detects permissive sites in the tomato pathogen *C. michiganensis* CASJ002. **A.** Whole genome analyses to detect permissive sites. Top: Presence and absence of genes encoded. Top-Middle: Functional IS elements predicted in the genome by ISEScan. Bottom-Middle: A 1000 bp scanning window calculation of Shannon entropy. Bottom: A 1000 bp scanning window of GC-content calculated along the genome with candidate permissive sites highlighted in blue. **B.** Subplots of predicted permissive sites ranked by GC-content and Shannon entropy. Left to right: top two ranked sites (HR1 and HR2) and last rank site (HR10).

**Figure 4:**
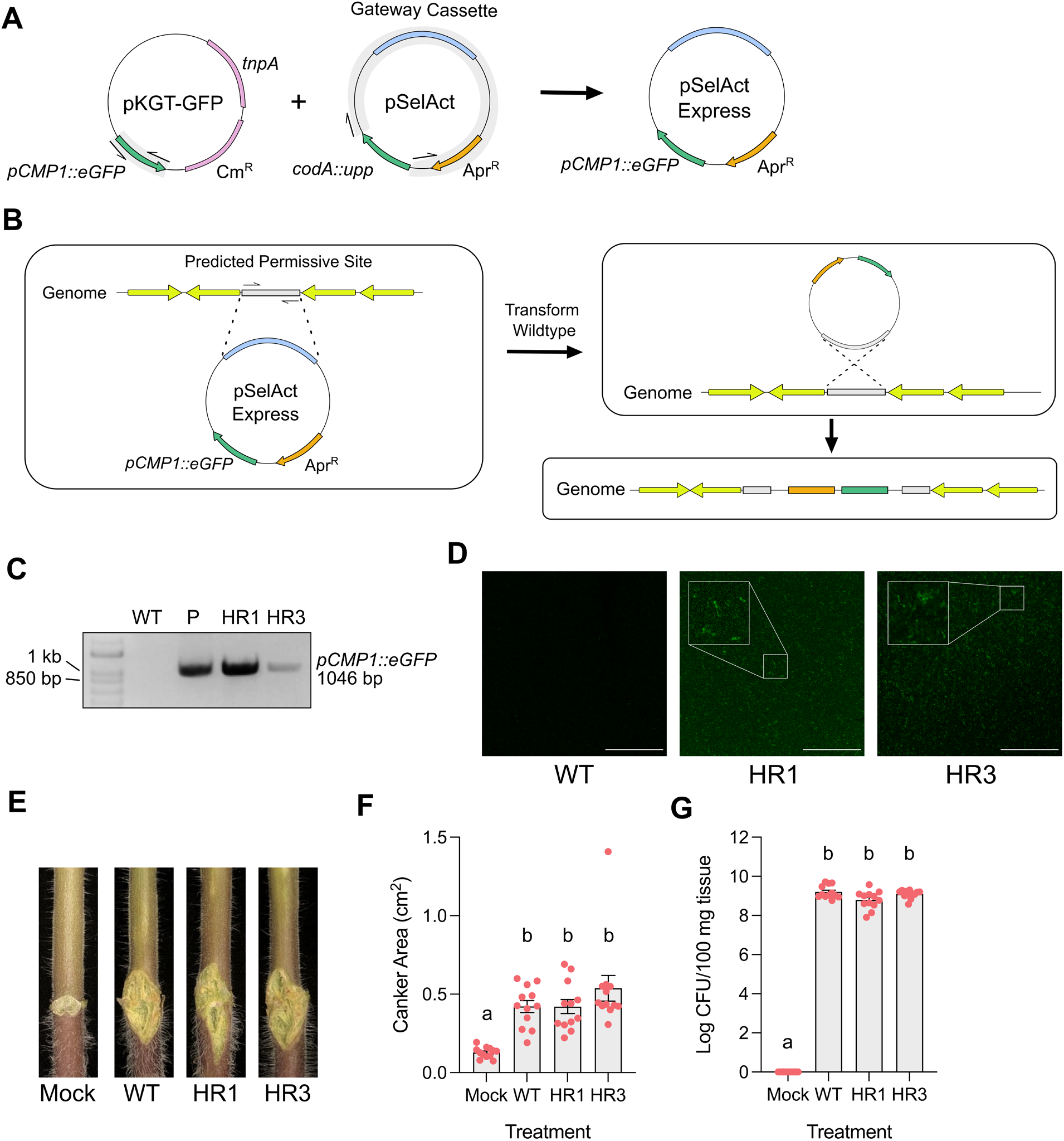
eGFP expression in two *Clavibacter* permissR target regions using an integrative plasmid. **A.** pSelAct-Express was developed by replacing the *codA::upp* counterselection cassette in pSelAct-KO with eGFP under a phage promoter from pKGT-GFP. **B.** Left: Amplification of a predicted permissive site from permissR into pSelAct-Express via Gateway or homology-based cloning. Right: Transformation with the modified non-replicating vector into the wildtype strain causes recombination into the genome at the predicted permissive site. **C.** PCR-based verification of the plasmid insertion via amplification of *pCMP1::eGFP*. WT; Genomic DNA from CASJ002, P: Plasmid DNA from *E. coli* carrying the pSelAct-Express; HR1: Genomic DNA from CASJ002 integration variant at hit region 1; HR3: Genomic DNA from CASJ002 integration variant at hit region 3. **D.** Confocal microcopy of eGFP expression in *C. michiganensis* CASJ002 (WT) and transformed integration variants (HR1 and HR3). White bar represents 40 μm in length. **E.** Representative tomato canker symptoms after inoculation with mock (water), wildtype (WT) CASJ002 and integrative variants HR1 and HR3. Stems were stabbed with a needle and inoculated with 5 ul of inoculum (either water or bacteria at OD_600_ equal to 1). Images were taken at 14 dpi. Four plants per treatment per experiment were conducted with the experiment repeated three independent times. **F.** Quantification of canker area from Figure E. Measurements were normalized with a one cm ruler (not shown). A one-way ANOVA and a post-means Tukey’s multiple comparisons test was computed. **G.** Bacterial titers in stem tissue 14 dpi at one cm above the site of inoculation from tomatoes in (E and F). A one-way ANOVA and a post-means Tukey’s multiple comparisons test was computed.

Bacterial assembly and annotation quality should be independently assessed before running permissR. Since the pipeline relies on space between coding information, permissR will not predict sites on bacterial genomes that are too fragmented. While permissR itself does not assess annotation quality, results are dependent on a reliable annotation. Common measurements of annotation quality include BUSCO scores, the number coding genes, and the number of hypothetical proteins. If the BUSCO scores are low or if the number of coding genes or hypothetical proteins becomes unreasonable compared to related taxa, then the genome should be improved before proceeding (Richardson and Watson 2013).

To test permissR, we used the *C. michiganensis* strain CASJ002 to predict potential sites. permissR predicted 10 candidate sites in the CASJ002 genome (Fig. 3A; Supplemental Table S3). Of the 10 candidates, one had a homologous region within the genome, which happened to be the second top predicted site (Fig. 3B). The GC-content of sites ranged from 55 to 75%. Additional *Clavibacter* genomes were tested against the pipeline to see if sites could be detected in other species (Supplemental Table S4). The number of predicted sites differed across *Clavibacter* strains and species including bacteria outside of the *Clavibacter* genus (Supplemental Table S4). This pipeline could, in theory, be used to help other researchers design integrated plasmids for their system of interest as permissR is not genera dependent.

### GFP expression in the *Clavibacter* chromosome using pSelAct-Express

Next, we sought to experimentally validate two candidate sites identified by permissR. First, we generated an integrative plasmid, pSelAct-Express containing GFP. pSelAct-Express was built from the pSelAct-KO vector by replacing the *codA::upp* counterselection cassette with the expression cassette from pKGT-GFP, which includes the phage promoter pCMP1 and eGFP (Fig. 4A; Chalupowicz et al. 2012; Tancos et al. 2013). We developed primers with homologous ends to amplify the pSelAct fragment and pCMP1::eGFP casette and cloned them together using In-Fusion. Similar to pSelAct-KO, to use pSelAct-Express the Gateway cassette is replaced with a ~1.5 kB region amplified from within one of the predicted permissive sites to facilitate recombination.

For testing the system in *C. michiganensis* CASJ002, ~1.5 kB regions within two permissive regions, hit region 1 (HR1) and 3 (HR3), were cloned into pSelAct-Express (Fig. 4B; Supplemental Table S5). Each construct was separately transformed into CASJ002 and construct integration was verified by PCR (Fig. 4B and C). Expression of eGFP was confirmed by confocal microscopy using wildtype CASJ002 as a negative control (Fig. 4D). To verify plasmid integration did not affect disease development and bacterial titers, three-week-old tomato cultivar M82 plants were stab inoculated. There were no significant differences between wildtype and the integrated variants when measuring the area of stem canker, a signature symptom in *C. michiganensis* disease development in tomato (Fig. 4E and F). Similarly, no significant differences were detected in bacterial titer 1-cm above the site of inoculation (Fig. 4G). While we did not observe alterations in bacterial fitness or disease development upon plasmid integration in HR1 or HR3, we recommend testing several candidate regions as there is always a potential for unpredictable effects. These results demonstrate permissR can be used to discover promising sites for plasmid integration using pSelAct-Express. We were able to identify two chromosomal regions for integrative expression in *C. michiganensis* that do not affect bacterial virulence.

## Conclusion

The tools optimized and developed in this work, pSelAct-KO, permissR, and pSelAct-Express, provides flexibility for genetic manipulation of *Clavibacter* bacteria. Molecular tools that enable manipulation for genotype-phenotype studies have greatly advanced our understanding of these organisms, and we conclude that the tools present here will open the door to investigating *Clavibacter*. Designed with flexibility in mind, we think there is a potential to be adapted to other orphan systems beyond bacteria in the *Clavibacter* genus.

## Acknowledgements

This research was supported by NIH 1R35GM136402 to GC and USDA-NIFA 2021-67034-35049 to DMS. We would like to thank Dr. Jeff Chang from Oregon State University for providing the pSelAct-KO vector and Qingyang Lyu for Sanger sequencing it. We would also like to thank MIGS for Illumina sequencing as well as Nick Carleson and Zarchary Foster for the suggestion of turning permissR into an R package. Finally, we would like to thank the members of the Coaker lab for their thoughtful discussion and reading of the manuscript.

## Author Contributions

DMS and GC designed the study. DMS and AT performed the experiments. DMS performed the bioinformatic analysis. All authors were involved in writing and editing the manuscript.

## Data Availability

Paired-end Illumina reads of *C. michiganensis* CASJ002Δ*chpE-ppaC* were submitted to Figshare (doi: 10.6084/m9.figshare.14810322). The scripts necessary for analyzing the data can be found in the GitHub repository (https://github.com/DanielleMStevens/pSelAct_KO_Clavibacter). The R package permissR can be found in the Github repository (https://github.com/DanielleMStevens/permissR) including installation and usage instructions.

## Materials and Methods

### Plasmids, Stains, and Culture Conditions

*C. michiganensis* strains were grown in tryptone broth with yeast (TBY) (Kirchner et al. 2001). *Clavibacter* transformants were selected on TBY rich media supplemented with 200 μg/mL apramycin (Apr) and screened on TBY with only 100 μg/mL apramycin (Alfa Aesar, Haverhill, MA, USA). *Escherichia coli* was grown in Luria Broth (LB) at 37°C. All strains were grown under standard temperatures listed in Supplemental Table S1 except when selecting for second recombinants. This process requires growth on a minimal media, M9, supplemented with 100 μg/mL of 5-florocystosine (5-FC) and grown at 24°C as listed in Supplemental Table S1 (Stork et al. 2008) (VWR, Radnor, PA, USA). A 10 mg/mL stock solution of 5-FC was prepared in distilled water and filtered sterilized. Apramycin was prepared similarly at a stock concentration of 100 mg/mL.

### Minimum Inhibitory Concentration of Apramycin in *Clavibacter*

Initial starter cultures were grown overnight at 28°C with shaking at 200 rpm in 5 mL of TBY media. Cultures were spun down at max speed for 2 min, washed with and resuspended in sterilize water. The optical density (OD) was taken at 600 nm and adjusted to a final concentration of 0.01 in TBY with different antibiotic concentrations. Eight 200 ul wells were used for each antibiotic concentration in a clear 96 well plate (Beckman Coulter, Pasadena, CA, USA) and incubated at 28°C with shaking at 200 rpm. OD_600_ measurements were taken at 0 and 18 hrs. Experiments were repeated a minimum three times.

### pSelAct-KO and pSelAct-Express

For generation of the multi-gene deletion plasmid, genomic DNA was extracted from 5 mL cultures of *C. michiganensis* CASJ002 grown overnight at 28°C with shaking at 200 rpm using a similar protocol of extraction (Murrary and Thompson 1980). ~1.5 kb regions flanking clustered genes of interest were amplified using PCR. It is important to select the position of the knockout carefully to avoid deletion into nearby protein coding genes to prevent unwanted polar effects. The reaction mixture for PCRs was as follows: 1x iProof GC Buffer, 0.2 μM of each primer, 50 to 200 ng of genomic DNA, 2% DMSO, 0.2 μM of DNTPs, and double distilled water to a final volume of 20 μl. Extension times were 15 to 30 seconds per kilobase of DNA amplified. Primer sequences, amplicon size, and optimized annealing temperatures for each primer pair are listed in Supplemental Table S5. The backbone of pSelAct-KO was linearized using Phusion polymerase with GC-buffer and required no more than 15 ng/ul of initial plasmid DNA. Amplicons were gel purified using the Zymo gel extraction kit according to the manufacturer’s instructions (Irvine, CA, USA). The knockout construct was developed through In-Fusion using the purified amplicons based on the manufacturer’s instructions with the insert to vector ratio of 2:1 (Takara Bio, Mountain View, CA, USA) and transformed into *E. coli* DH5α. The resulting plasmid insertions were amplified and confirmed via Sanger sequencing.

For generation of pSelAct-Express, pKGT-GFP plasmid DNA was extracted from *E. coli* using a QIAprep Spin Miniprep Kit according to manufacturer’s instructions (Qiagen, Germany, Hilden, Germany). The phage promoter eGFP expression cassette (*pCMP1::eGFP*) was amplified via PCR (Supplemental Table S5). The pSelAct-KO plasmid was linearized without *codA::upp* using similar conditions as stated above. Similar to generating the knockout construct (see above), the plasmid was generated using In-Fusion, transformed into *E. coli* DH5α, screened via PCR, and confirmed via Sanger Sequencing.

For cloning into hit regions from permissR predictions in *C. michiganensis* strain CASJ002, primers were designed from output sequence to amplify regions ~1.4 to 1.5 kB in size. HR1 and HR3 were amplified from genomic DNA and cloned to replace the Gateway cassette via In-Fusion (Supplemental Table S3 and S5). Positive clones were confirmed as described above.

### Transformation and Selection

*Clavibacter* competent cells were prepared and transformed as previously described (Kitchner et al. 2001). Briefly, cells were transformed with 100 to 200 ng of plasmid DNA and plated on TBY with 200 ug/mL apramycin. Colonies were picked and re-streaked onto TBY with 100 ug/mL of apramycin. Colonies were cultured overnight in TBY with 100 ug/mL apramycin for genomic DNA extraction (Murrary and Thompson 1980).

### Selection for 2^nd^ Recombinant

Positive merodiploids were grown overnight at their respective temperature and media. The culture was centrifuged at max speed, pellet resuspended in 1 mL sterile water. Ten-fold dilutions from 10^-1^ to 10^-7^ were made and 100 ul from 10^-6^ and 10^-7^ dilutions were plated onto mM9 medium supplemented with 100 ug/mL of 5-FC. Plates were incubated at their indicative lower temperature, 24°C, optimized for counter-selection (Supplemental Table 1) for 9 to 14 days. To screen quickly for recombination events, colonies were replated onto their indicated rich media with and without apramycin selection. Positive colonies, which only grew on the rich media, were additionally screened via PCR using both span and gene specific primers for a true knockout and not a recombinant that has returned to wildtype state.

### Bioinformatic Analyses

*C. michiganensis* CASJ002Δ*chpE-ppaC* genomic DNA was prepared from 5 mL overnight cultures (Murrary and Thompson 1980). The Microbial Genome Sequencing Center (MIGS) library prepped the DNA for paired end read sequencing via an Illumina platform. Raw paired end reads were check for quality and any contamination using fastqc (v0.11.9) and the sendsketch.sh script from bbtools, respectively. Since contamination was present, genomes closest to the top one to two hits from the sendsketch.sh output were downloaded using the bioinf_tools package (v1.8.17) (Lee 2018). Reads were then mapped to the genome of the *Clavibacter* reference and contaminants and binned to their respective hit using bbtools’ bbsplit.sh script. Successful binning was confirmed using sourmash (v3.5.0) based on a kmer size of 31 (Brown and Irber 2016). Merged reads were reformed via reformate.sh from bbtools and trimmed using trimmomatic (v0.39). Trimmed reads were *de novo* assembled via spades (v3.14.1) with default parameters for short read assembly (Bankevich et al. 2012). Contigs were scaffolded using medusa based on the reference wildtype genome and oriented based on the reference using contiguator2 (Galardini et al. 2011; Bosi et al. 2015). The scaffolded contigs and the region of interest were mapped to their associate reference via minimap2 (v2.17-r941) and fastANI (v1.32.0) and structural changes were assessed using a custom R script which uses the R package pafr (0.0.2) and genoplotR (0.8.11) (Charif and Lobry 1007; Li 2018; Jain et al. 2018). Detailed methods and scripts can be found at (https://github.com/DanielleMStevens/pSelAct_KO_Clavibacter).

### The R package permissR

permissR was written in the R language (R Core Team 2019) and requires only two input files, at minimum, a Genbank file (gbk or gbff file formate) and a Fasta file. While the package does not require any external programs to run, it is recommended running and including the output from ISEScan, an annotation-independent software to find IS elements in bacterial genomes to avoid the potential of cloning into an functional IS element that may cause unwanted gene movement (Xie and Tang 2017). To run ISEScan, we recommend using the package management software bioconda.

Detailed installation and usage can be found in the permissR GitHub repository (https://github.com/DanielleMStevens/permissR). Briefly, installation of the package requires installing the *devtools* R package and uses devtools::install_github(“DanielleMStevens/permissR”). To run, calling the function permissR in the package on the R console, the user can provide the path (relative or absolute) for the Genbank file and then Fasta file of interest or leave the function empty, which will cause a GUI interface to appear to allow the user to select each file. The function will ask the name of the strain, which is used to label the output files as well as if there are any outputs for ISEScan.

### Plant growth and pathogenicity assays

The tomato cultivar M82 (*Solanum lycopersicum* cv. M82) was used for all assays. Tomato plants were grown at 25°C in a growth chamber under 16 hrs light:dark conditions with 50% humidity and at least 200 lumens. Plants were grown for 3 weeks when at least two true leaves had fully emerged, pricked between the two cotyledons on the stem using a sterilized needle, and 5 ul of bacterial inoculum or water (mock) was dropped into the wound. The inoculum was prepared by initially streaking out *Clavibacter* stocks four days before inoculation on either TBY or TBY plates with apramycin. The day of inoculation, bacteria was suspended from the plate in sterile water and optical density at 600 nm was adjusted to 1. Symptoms were photographed at 14 dpi and canker was measured using ImageJ based on a 1 cm ruler to normalize.

To determine bacterial titers in the tomato stem, stem segments were cut surrounding the inoculation site at 14 dpi. Stem pieces were surfaced sterilized in 70% ethanol for 10 to 15 seconds followed by suspension in sterile water for 10 to 15 seconds. Sections were taken 1 cm above inoculation site. Tissue was weighed at 100 mg, suspended in sterile water, and ground. Sterile dilutions of the ground tissue were plated on D2 medium supplemented with 20 mg of cycloheximide and incubated for five to six days at 28°C (Thapa et al. 2017).

### Confocal microscopy

Before imaging, *Clavibacter* stocks were streaked out onto either TBY or TBY plates with apramycin and grown at 28°C. Bacteria was smeared onto glass slide, 2 μl of sterile water were pipetted on, and a glass cover slip was added on top. Confocal images were taken with a Leica SP8 TCS with a 63x oil objective, laser power set to 6%, and excitation and emission wavelengths set to 488 nm and 509 nm, respectively. Fiji with the Bio-Formats Importer was used to adjust the brightness and contrast of the images.

### Statistical analyses

All raw data was plotted and error bars in plots represent standard error of mean (SEM). Data across independent replicates was colligated and a one-way ANOVA and a post-means Tukey’s multiple comparisons test was computed using Graphpad Prism 9 software. Significant differences between groups include P values equal to or less than 0.001. Outputs of statistical analyses from Figure 2B, Figure 4F, and Figure 4G can be found in Supplemental Table 6, 7, and 8, respectively.

---

**Supplemental Figure 1:**
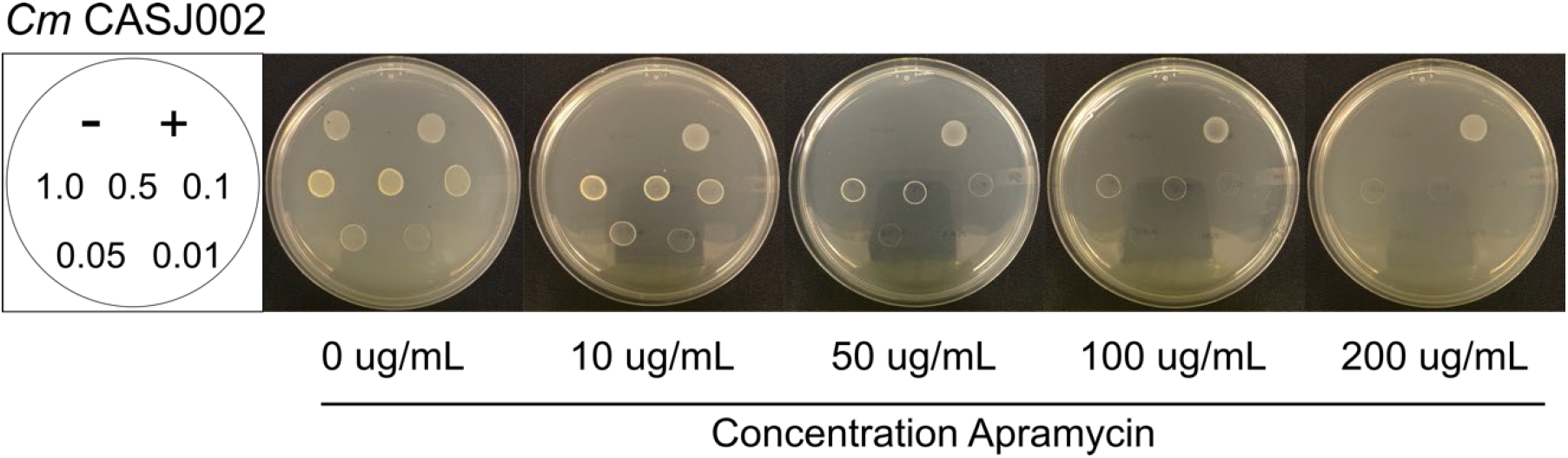
Concentration of apramycin required to restrict *C. michiganensis* on agar plates. Left: Schematic for plating. The negative sign represents the negative *E. coli* control. The positive sign represents the positive control, the pSelAct-KO vector in *E. coli*. Both controls are tested at an OD_600_ of 1. The numbers 1.0 through 0.01 are the tested optical densities (600 nm) of *C. michiganensis* isolate CASJ002. Right: Apramycin concentrations in TBY media.

**Supplemental Figure 2:**
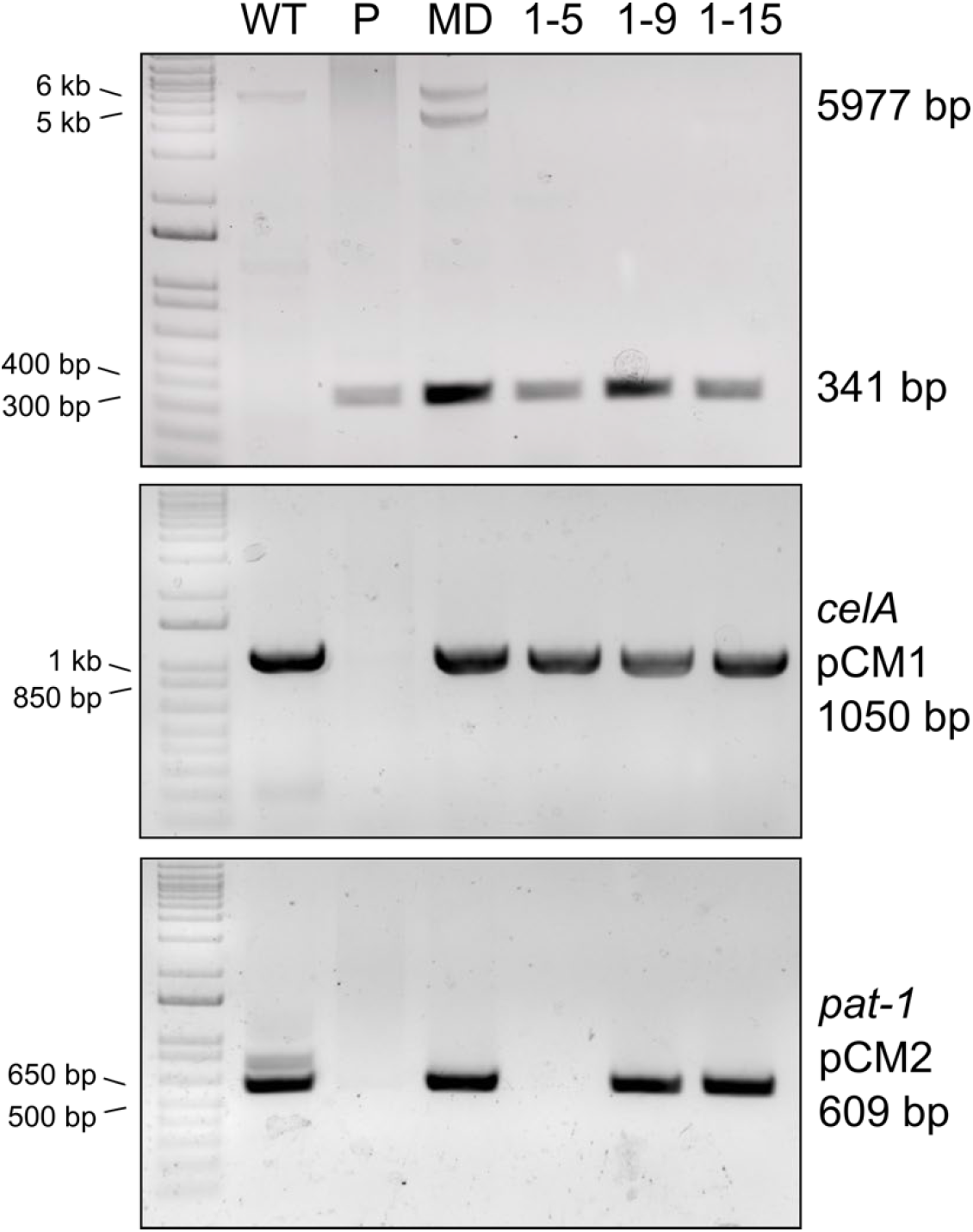
Counterselection can lead to loss of native plasmids. PCR-based verification of native plasmids pCM1 and pCM2 on a 1% TAE agarose gel. Top: Primers which bind to the flanking regions and span the deleted region. Middle: Gene specific primers for *celA* (indicates presence of pCM1). Bottom: Gene specific primers for *pat-1* (indicates presence of pCM2). Lanes are products from the reactions including: WT; Genomic DNA from CASJ002, P: Plasmid DNA from *E. coli* carrying the pSelAct_CASJ002_chpE-ppaC_KO; MD: Genomic DNA from the merodiploid; 1-5, 1-9, and 1-15: Genomic DNA from three CASJ002Δ*chpE-ppaC* recombinant colonies. The estimated sizes of the PCR products are depicted.

**Supplemental Figure 3:**
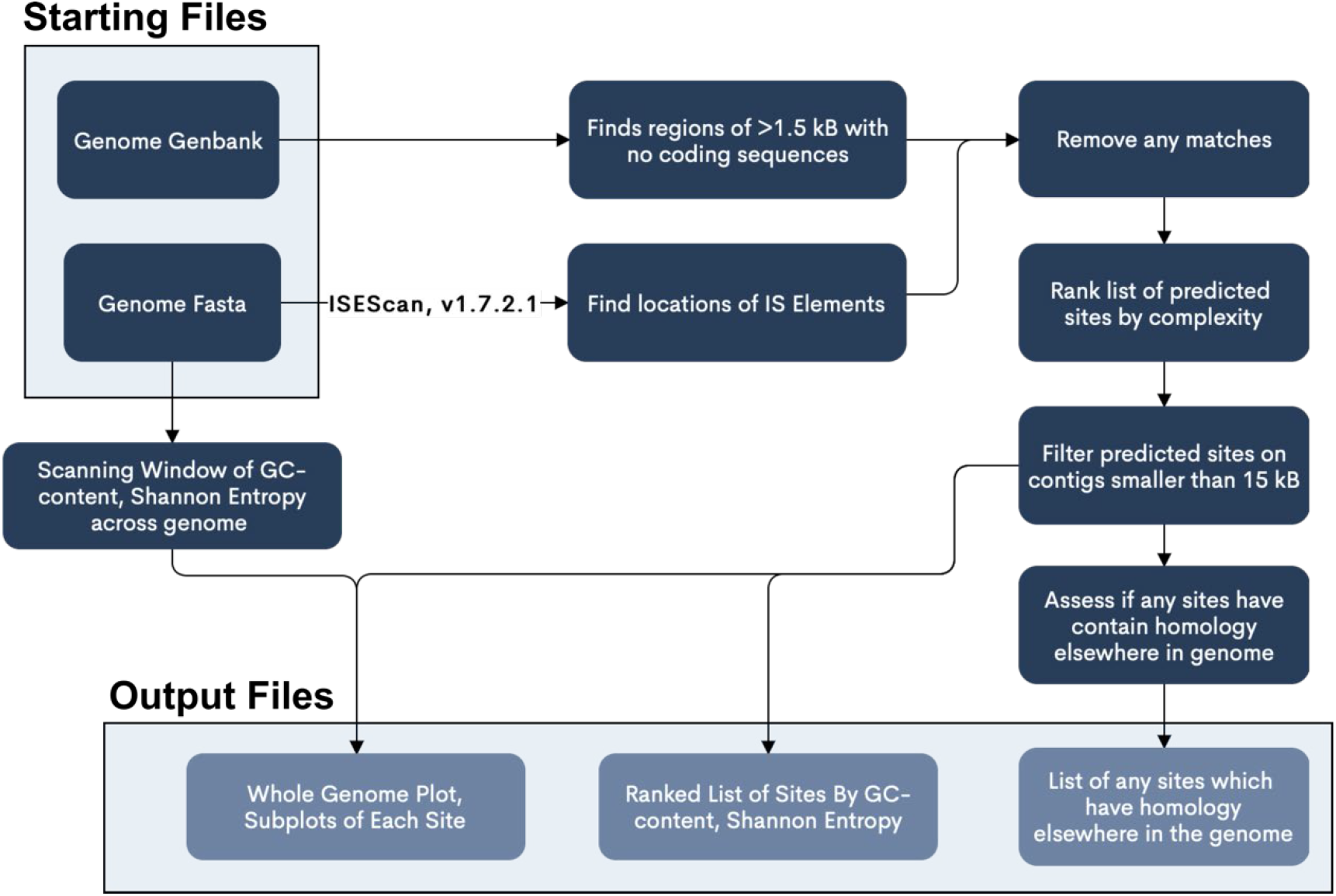
Pipeline for the R package permissR to predict permissive sites in bacterial genomes. The pipeline uses whole genome Genbank and Fasta files to identify sites which are at least 1.5 kB in length (for specific recombination), that contain no coding elements, no mobile elements (which may trigger structural changes), and are ranked by complexity to make cloning reasonable (i.e. too high GC-content).

